# Unlocking the Bile Acid Universe: Advanced Workflows and a Multidimensional Library of 280 Unique Species

**DOI:** 10.64898/2026.02.18.706700

**Authors:** Guozhi Zhang, Emily C. Vincent, Sadie M. Disselkoen, James N. Dodds, Quentin DuVal-Smith, Abubaker Patan, Ipsita Mohanty, Victoria Deleray, Jian Zhang, Paul A. Thiessen, Evan E. Bolton, Emma L. Schymanski, Pieter C. Dorrestein, Casey M. Theriot, Erin S. Baker

## Abstract

Microbes and bile acids are tightly intertwined, especially in the gut. While the liver produces primary bile acids from cholesterol, gut bacteria transform these into diverse secondary forms which act as powerful signaling molecules, influencing host metabolism and immune function. Since bile acid changes are increasingly linked to health and disease, their accurate measurement in the gut and circulation is essential. Analytical evaluations, however, remain challenging as many bile acids co-elute in liquid chromatography (LC), share identical precursor masses in mass spectrometry (MS), and produce similar tandem mass spectrometry (MS/MS) spectra. As a result, conventional LC-MS/MS workflows struggle to differentiate bile acids, motivating the addition of orthogonal separations such as ion mobility spectrometry (IMS). Here, we assess optimal bile acid extraction parameters for stool, serum, and plasma; compare LC conditions; and assess electrospray ionization performance across polarities. Additionally, we created a multidimensional reference library containing LC retention times, IMS collision cross section values, and accurate precursor masses for 280 unique bile acids (264 endogenous and 16 deuterium-labeled species) including unconjugated, host-conjugated, and microbially conjugated bile acids. This multidimensional library empowers bile acid identification in complex samples and enables a more comprehensive exploration of their biological roles and disease associations.

## Introduction

Bile acids are structurally diverse bioactive molecules crucial for lipid digestion, cholesterol metabolism, and host-microbiota interactions^1–5^. Beyond their well-established functions in dietary fat emulsification, bile acids act as potent signaling molecules that regulate metabolic pathways, immune responses, and gut microbial composition^1, 5–7^. Disrupted bile acid metabolism is therefore implicated in metabolic syndrome, inflammatory bowel disease (IBD), and microbiota-associated dysbiosis, highlighting their potential as both diagnostic biomarkers and therapeutic targets^8–14^. Recently, it was found that beyond glycine- and taurine-conjugated bile acids, gut bacteria produce microbially conjugated bile acids (MCBAs), linking molecules such as amino acids and novel neuroactive conjugates such as gamma-aminobutyric acid (GABA) to bile acids^15–20^. These microbial modifications increase bile acid functional diversity and influence host neurotransmitter signaling, gut-brain axis interactions, and immune modulation^12, 21, 22^. Emerging evidence further links MCBA modifications to neurological outcomes, with altered conjugate profiles observed in extremely premature infants with brain injury^23^, reinforcing their importance. Given their biological specificity and structural diversity, accurate annotation and quantification of the many possible bile acids, particularly MCBAs, is essential for understanding their distinct biological roles in health and disease.

Despite their biological importance, identifying and evaluating bile acids remains analytically challenging due to their extensive isomeric diversity and microbial modifications^21^. While widely used for bile acid profiling, conventional liquid chromatography and tandem mass spectrometry (LC-MS/MS) methods often struggle to resolve isomeric species due to LC co-elution and overlapping mass-to-charge (*m/z*) ratios^2, 24–30^. Reliance on MS/MS fragmentation patterns alone is also typically inadequate for structural annotation since the conserved steroid core structures of bile acids often produce nearly identical fragmentation spectra, limiting differentiation of positional and stereoisomeric bile acid isomers^31^. Furthermore, a recent study reported that 176 unique MCBAs corresponded to only 42 distinct *m/z* values, highlighting limitations of mass-based separations alone^32^. Coupling ion mobility spectrometry (IMS) with LC and MS separations (LC-IMS-MS) has emerged as a promising analytical enhancement for bile acid analyses with previous studies showing improved bile acid isomer differentiation^33–44^. However, many collision cross section (CCS) values are missing for bile acids, particularly for MCBAs, limiting confident annotation in complex samples and motivating expansion of multidimensional reference measurements^21, 33^. In many cases, as this work shows, even the bile acid structures themselves are absent from databases used for metabolite annotation.

Beyond difficulties in bile acid analytical separation and annotation, the systematic evaluation of analytical method choices remains limited including: (1) extraction methods that balance recovery and feasibility across matrices and (2) LC conditions that separate bile acid species relevant to a given study^25, 45^. Currently, bile acid extraction workflows differ substantially by matrix. For blood matrices, acetonitrile and methanol are widely used to extract bile acids since they efficiently precipitate proteins.^2^ In contrast, fecal approaches span isopropanol-based protocols optimized for oxo-bile acids at 37°C to emerging dried fecal spot (DFS) methods that enable remote sample collection^46^. Many protocols are also labor-intensive and difficult to scale, and the limited number of head-to-head comparisons makes it unclear which parameters most strongly drive bile acid recovery.^47, 48^ Furthermore, once bile acids are extracted, it is often difficult to know which LC methods will distinguish the bile acids of interest. To address these gaps, we systematically evaluated extraction solvent changes for blood and stool samples, evaluated chromatographic conditions and their trade-offs, and characterized bile acid ion formation in positive and negative MS source conditions.

Using our LC-IMS-MS workflow, we created a publicly available multidimensional bile acid reference library encompassing bile acid structures, retention times across three LC methods, experimental CCS values in positive and negative mode, and accurate masses for 280 unique bile acids. The library therefore contains 343 deprotonated features in negative ionization mode and 665 features (including sodiated, ammoniated and protonated species) in positive mode (**Tables S1-S2**). We expect this resource to enhance bile acid identification and facilitate bile acid profiling studies across host and microbiome metabolism.

## Methods

### Standard Preparation

#### Bile Acid Standard Preparation

Bile acid standards and mixtures were purchased from Cayman Chemical (Ann Arbor, MI) and Cambridge Isotope Laboratories (Tewksbury, MA) **(Supplemental Information)**. In addition to the Cayman and CIL standard mixtures, 22 individual unlabeled standards (^12^C) were purchased from Cayman Chemical and synthetic bile acids were received from the Dorrestein lab and most can now be obtained from BileOmix, while others are part of a legacy collection from the late Alan Hoffmann and provided by Lee Hagey^8, 49^. The standards were then used to populate the LC-IMS-MS reference library and support bile acid annotation (**Tables S1-S3**). Individual unlabeled bile acid standards from Cayman were supplied in variable quantities, and accurately weighing sub-milligram amounts is prone to large errors. Therefore, each individual standard was reconstituted in 500 μL methanol to generate a stock standard in solution, vortex-mixed until fully dissolved, and stored at −20°C to minimize degradation. Because bile acids differ substantially in solubility and electrospray ionization (ESI) efficiency, the concentrations required to achieve comparable on-column signal vary by analyte. Accordingly, individual standards were prepared across a broad concentration range (0.5-100 mg/mL) to obtain measurable signal for each compound under the tested LC-IMS-MS conditions. The Cayman Bile Acids MaxSpec® Discovery Mixture (18 bile acids, catalog number 33505) was also utilized for LC method optimization and received in methanol and used without further preparation.

#### Internal Standard Preparation

A ^13^C-labeled bile acid internal standard mixture was used as an extraction process control to monitor extraction consistency, instrument stability, and confirm successful extraction. The mixture was prepared by combining two CIL bile acid mixes: labeled conjugated bile acid mix (MSK-BA1-1) and a labeled unconjugated bile acid mix (MSK-BA2-1). Each vial was reconstituted in 1000 μL of a 1:1 methanol/water mixture, and the solutions were combined to yield a 2000 μL internal standard mix with an approximate concentration of 50 μM per bile acid (0.02 mg/mL MSK-BA1-1 and 0.025 mg/mL MSK-BA2-1). This internal standard was spiked into samples prior to extraction to monitor instrumental/extraction performance but was not used for signal normalization or quantification in the analyses reported here.

### Bile Acid Extraction from Stool

The extraction procedure followed previously established protocols^50^. A step-by-step SOP for solid matrix extraction is provided in **Supplemental Information**. Approximately 50 mg of a control human stool sample^50^was aliquoted into eight 2 mL microtubes containing 2.38 mm metal beads (Omni International, Kennesaw, GA, USA; catalog number 19-620) to compare four different extraction solvent conditions. Due to sample limitations, each extraction condition was performed in duplicate to ensure reproducibility. The exact stool conditions and masses used for each replicate are provided in **Tables S**4**-S**5, and the extraction conditions are noted below.

#### Pre-extraction Buffer Preparation

Four different HPLC grade organic solvents were evaluated for stool extraction: ethanol (Sigma-Aldrich, St. Louis, MO, catalog number 226557), methanol (Fisher Scientific, catalog number 67-56-1), acetonitrile (Fisher Scientific, Hampton, NH, catalog number 75-05-8), and a 1:1 (v/v) mixture of acetonitrile (ACN) and methanol (MeOH) **(Table S4)**. The four solvents were selected based on their effectiveness in extracting bile acids of varying hydrophobicity, as supported by previous studies^50^. Each pre-extraction buffer was prepared by diluting a 100 mM potassium phosphate monobasic buffer (Sigma-Aldrich, catalog number 7778-77-0) 1:5 (v/v) with water to achieve a 20 mM aqueous phosphate solution. Pre-extraction buffer was then prepared by combining 60 mL of the 20 mM phosphate solution with 340 mL designated organic solvent to obtain a 15:85 (v/v) aqueous:organic ratio (total volume 400 mL) and a final phosphate concentration of 3 mM. For the 1:1 acetonitrile:methanol condition, the 340 mL organic fraction is comprised of 170 mL acetonitrile and 170 mL methanol. Each pre-extraction buffer was added to the stool samples at a ratio of 8 μL per mg of sample (*e.g.,* 400 μL for 50 mg) to account for minor variations in sample mass.

#### Sample Extraction and Processing

Briefly, prior to extraction, a fixed volume of the ^13^C-labeled bile acid internal standard mix was added to each tube. Samples in pre-extraction buffer were homogenized at room temperature using a Fisherbrand™ Bead Mill 24 Homogenizer (Fisher Scientific, catalog number 15-340-163) at 2.1 m/s for 2 min in a single cycle to ensure complete disruption. The homogenized samples were centrifuged at 13,000 × g for 10 min at 4°C. The resulting supernatant from each extraction condition was divided into three fractions: 100 μL was mixed 1:1 with methanol, 50 μL was mixed 1:4 with methanol, and 50 μL was mixed 1:4 with the same organic solvent (without phosphate) used in the corresponding pre-extraction step. Each mixture was vortexed at 600 rpm for 20 min at room temperature to ensure thorough extraction. The extracted samples were then filtered using MilliporeSigma PTFE membrane spin filters (0.2 μm pore size, catalog number UFC30LG25), and a 60 μL aliquot of the filtrate was transferred to Agilent LC vials (Agilent Technologies, catalog number 5188-6591) for analysis.

### Bile Acid Extraction from Serum and Plasma

Bile acids were extracted from human control serum and plasma samples, and NIST serum 909C and NIST plasma 1950 using a 1:1 (v/v) acetonitrile:methanol with a detailed supplemental SOP and **Table S6**. These extraction solvents were added at a 1:4 ratio to two different sample volumes of serum and plasma (200 μL and 50 μL) to compare high and low sample quantities, and performed in triplicate (n=3), resulting in 24 samples for the serum and plasma including NIST reference (*e.g.,* 4 matrices × 2 volumes × 3 replicates). Additionally, a 1:8 extraction ratio was also evaluated using 50 μL of the sample, but this was not replicated due to sample limitations.

#### Sample Extraction and Processing

A fixed volume of the ^13^C-labeled bile acid internal standard mix was added to each sample above. Samples were vortexed at 2,000 rpm for 5 min at room temperature, followed by sonication on ice for 15 min to enhance bile acid extraction efficiency. Samples were then incubated overnight at −20°C, allowing for protein precipitation. This step was included to ensure consistent extraction efficiency across different sample volumes. The following day, samples were centrifuged at 13,000 × g for 10 min at 4°C using an Eppendorf 5810R centrifuge. A 100 μL aliquot of the supernatant was transferred into a clean 2.0 mL Eppendorf tube (MilliporeSigma, catalog number 30123620) for further analysis.

### LC-IMS-MS Analysis

#### LC Separation

Injection volume (1-6 μL) was adjusted based on concentrations of each bile acid standard to maintain comparable peak shapes. For extracts, a fixed injection volume of 6 μL was used. All samples were injected into an Agilent 1290 UPLC coupled to an Agilent 6560 IMS-QTOF mass spectrometry instrument. All extracts and standards were evaluated in negative ESI mode, and positive mode was also used for bile acid standards for library building. Detailed mobile phase compositions, and gradient programs are provided in **S10** and **Table S7**.

The LC separation for the stool, serum, and plasma extracted samples were performed using a Restek Raptor C18 column (1.8 μm, 50 × 2.1 mm; Restek, catalog number 9304252), coupled with a ZORBAX Eclipse Plus C18 guard column (2.1 mm, 1.8 μm; Agilent Technologies, catalog number 821725-901). The guard column was used to protect the main analytical column from sample matrix components in biological samples and enhance reproducibility and separation consistency across analyses. The Restek column and neutral mobile phase were chosen based on their previous application in bile acid separations^32^, serving as a baseline condition for method comparisons.

For bile acid standards, three different LC methods were evaluated using both the Restek Raptor C18 column and the Waters ACQUITY Premier CSH C18 column (1.7 µm, 2.1 × 100 mm; Waters Corporation, catalog number 186009461). A total of three mobile phase sets, consisting of both neutral pH and acidic pH conditions, were tested. The column temperature was maintained at 60 °C for both columns with a flow rate of 0.5 mL/min for the Restek Raptor column and 0.4 mL/min for the CSH column.

#### Calibration

Prior to any LC-IMS-MS analyses, mass calibration and CCS calibration were performed using the Agilent ESI-L Low Tuning Mix (Santa Clara, CA; catalog number G1969-85000), ensuring accurate mass calibration and precise CCS measurements.

#### IMS-MS Acquisition

The Agilent 6560 IM-quadrupole time-of-flight (QTOF) MS instrument was used to acquire the IMS and MS data in the experiment. Following chromatographic separation and ESI, the ions traveled through a single-bore glass capillary, where they were focused into a high-pressure funnel and accumulated in a trap funnel. The ions were then pulsed into a 78 cm IMS drift tube containing 3.95 Torr of nitrogen drift gas, using Hadamard transform multiplexing^51^. A 4-bit pseudo-random pulsing sequence was applied for packet ejection from the trapping funnel, with a trap fill time of 3.9 ms and a release time of 150 μs. After traversing the drift tube, the ions were refocused in a rear ion funnel and analyzed using the QTOF over a range of 50-1700 *m/z*. The “.d” files were then collected containing information on the LC, IMS and MS values for each ion detected in the experiments. Details of the MS parameter can be found in **Table S8.**

#### CCS Calculation

Drift times were extracted from raw Agilent “.d” files using Agilent MassHunter IM-MS Browser 10.0. CCS values were then calculated using the single field^52^. For all reported analytes, *m/z* mass errors were under 10 ppm and CCS the percent relative standard deviation (%RSD) across replicates was below 0.3%.

### Skyline Targeted Bile Acid Analyses

Skyline 23.1 (MacCoss Lab, Seattle, WA, release date 9/24/23), was used for compiling retention time information from standards and targeted peak integration to annotate bile acids in the extraction samples. Bile acid standards and mixtures acquired using the three LC conditions were imported into Skyline and manually annotated based on the reference library developed in this study. Retention times for annotated targets were exported to Microsoft Excel and summarized in **Tables S1**-**S2**. Deuterated standards are listed in both the quantitation and deuterated sections of Tables S1 and S2 for ease of import into Skyline. For extraction analyses, stool, serum, and plasma “.d” data were imported into Skyline, and peaks for bile acids and MCBAs were integrated using the same target library list. Peak area under the curve (AUC) for each bile acid and MCBA, along with mass errors, was exported to Microsoft Excel for further analysis. Data points with mass errors exceeding 10 ppm were excluded from processing.

## Data Availability

All data from this study are publicly available. The raw mass spectrometry data (“.d” files) have been deposited in the MassIVE repository (https://massive.ucsd.edu) under accession number [MSV00096436]. The Skyline spectral library and processed results are available on Skyline Panorama (https://panoramaweb.org/PkMTb5.url). The bile acid library data, including LC-IMS-MS spectra, retention times, and CCS values, have been deposited in Zenodo (https://doi.org/10.5281/zenodo.18675023) and the structures and CCS values added to PubChem (see data subsets under the Baker Lab source https://pubchem.ncbi.nlm.nih.gov/source/25763 and CCS classification at https://pubchem.ncbi.nlm.nih.gov/classification/#hid=124). Protocols for bile acid extraction, including SOPs, are available in **Supplemental Information.**

## Results and Discussion

Evaluating analytical workflow choices is essential for confident bile acid annotation in complex matrices. Practical analytical choices that influence bile acid detectability and annotation in stool, serum and plasma samples were investigated further in this manuscript. Each optimization step is detailed below, including the construction of a multidimensional LC-IMS-MS reference library for all bile acid ions detected in both positive and negative ionization mode.

### Bile Acid Extraction Analyses

To support robust bile acid profiling across matrices, extraction is extremely important. Here we evaluated extraction workflows for (i) stool sample and (ii) serum and plasma, since matrix properties strongly influence recovery and detectability. **Figure 1** summarizes the experimental design and variables evaluated for each matrix, including extraction solvent composition and solvent-to-sample ratio. The stool, serum, and plasma extractions are detailed further below.

**Figure 1.**
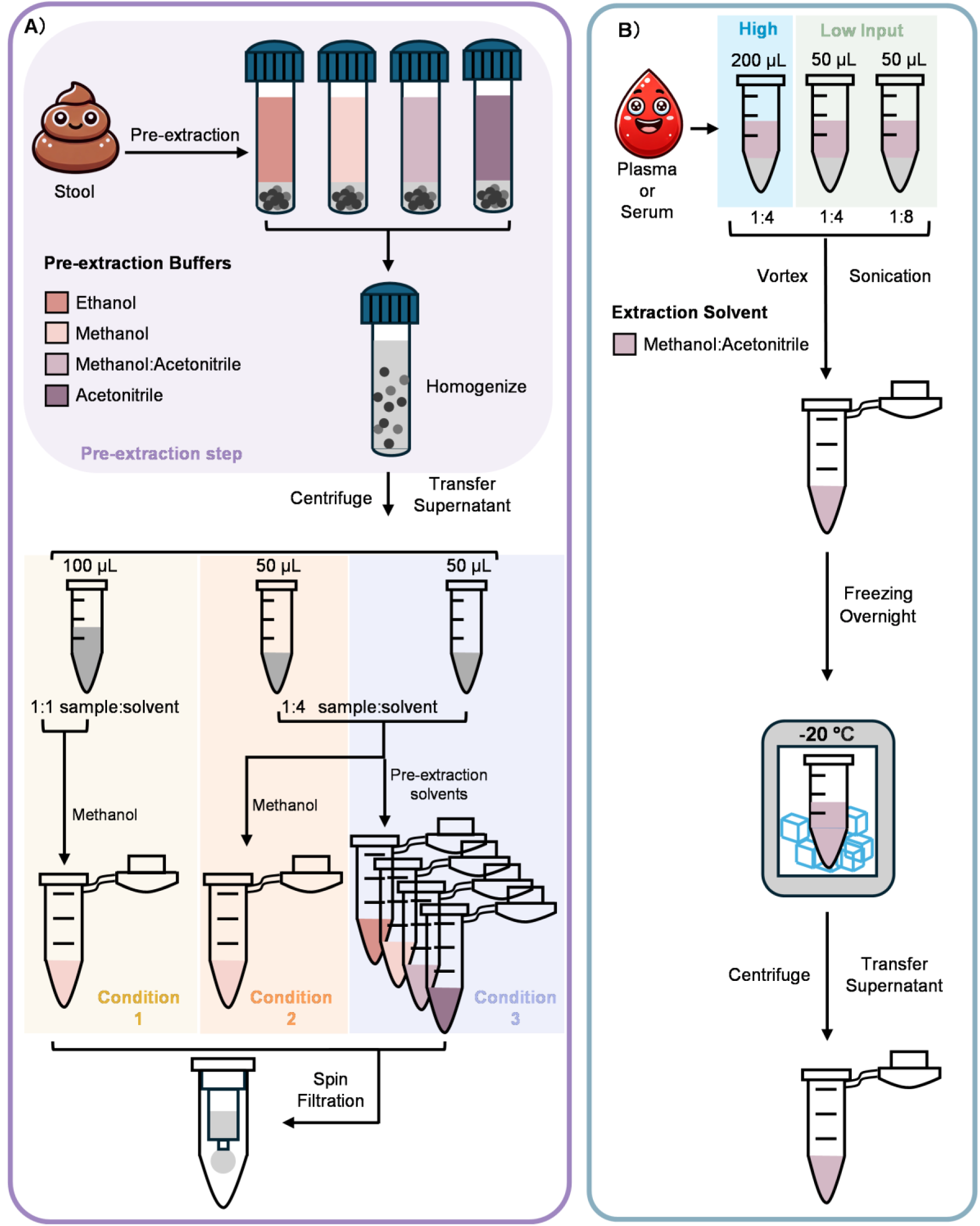
Extraction workflow and variables evaluated for stool and serum/plasma samples. **A)** Stool bile acids were extracted using a two-step workflow consisting of pre-extraction step (*purple*) with buffers (ethanol, methanol, acetonitrile, or 1:1 acetonitrile/methanol) and then followed by a second-step dilution condition. Three dilution conditions were evaluated, including (1) 1:1 (*yellow*) or (2)1:4 dilution in methanol (*green*), or 1:4 dilution in the matched pre-extraction solvent (*blue*)). **B)** Serum and plasma samples were extracted at high (200 μL) and low (50 μL) input volumes using a 1:1 acetonitrile/methanol solvent at a 1:4 sample-to-solvent ratio.

### Stool Bile Acid Extraction

Stool bile acids are commonly extracted using solvent-based homogenization/extraction with organic solvent systems, although protocols vary widely in solvent composition and sample-to-solvent handling, systematic comparisons across these practical choices are not consistently reported. Here, we separated the effects by varying (i) pre-extraction solvent buffer composition and (ii) the second-step dilution solvent, allowing us to evaluate how these practical choices affect extraction efficiency (**Figure 1A**). Briefly, approximately 50 mg of stool sample was extracted using a two-step workflow consisting of an initial solvent extraction (pre-extraction) followed by a second-step dilution using either methanol or the matched pre-extraction solvent. Four pre-extraction buffers (ethanol, methanol, acetonitrile, or 1:1 acetonitrile/methanol) were added directly to the raw stool in the pre-extraction step. In the second-step, three dilution conditions were then tested: Condition 1 - 1:1 dilution in methanol using 100 μL supernatant and 100 μL methanol (**Figure 2A**), Condition 2 - 1:4 dilution in methanol using 50 μL supernatant and 200 μL methanol (**Figure 2B**), and Condition 3 - 1:4 dilution in the matched pre-extraction solvent using 50 μL supernatant and 200 μL matched solvent (**Figure 2C**). We selected these volumes to reflect common high-throughput handling constraints and to test whether reduced carryover volume might result in the reduced detection or loss of certain bile acids. Specifically, the 1:4 conditions carry forward less first-step supernatant (50 μL vs 100 μL), which may reflect practical workflow choices.

**Figure 2.**
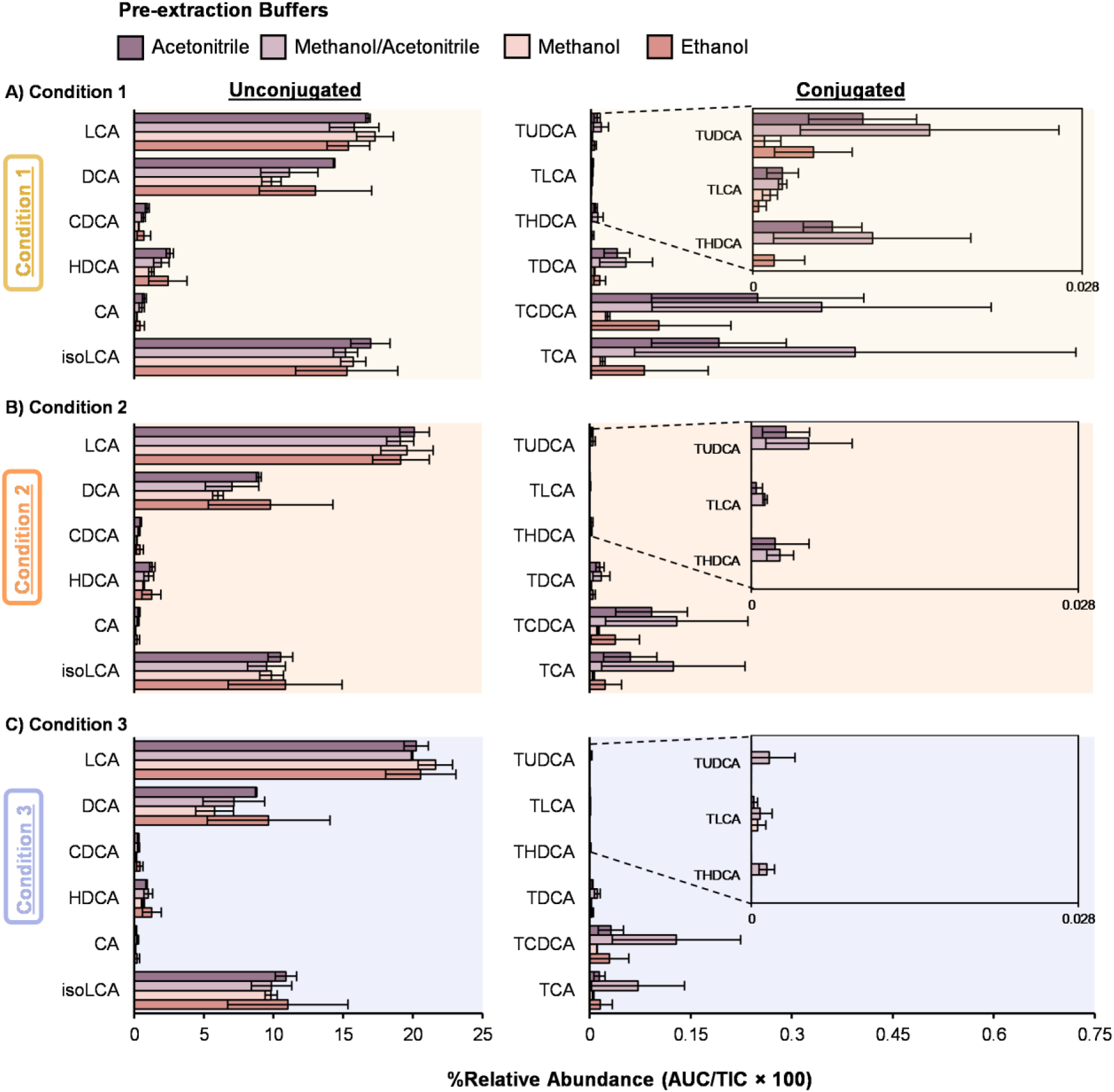
Extraction conditions on bile acid analysis from human stool samples. Bile acid extraction efficiency was evaluated using a two-step extraction workflow with different pre-extraction buffers (color-coded) and extraction ratios where: **A)** Condition 1 uses a 1:1 ratio with 100 μL of pre-extraction supernatant and 100 μL of methanol, **B)** Condition 2 has a 1:4 ratio with 50 μL of pre-extraction supernatant and 200 μL of methanol, and **C)** Condition 3 a 1:4 ratio with 50 μL of pre-extraction supernatant and 200 μL of matching organic solvent as pre-extraction solvent. A Skyline search against the full library detected 34 bile acids, and a representative subset of 12 are shown with unconjugated (*left*) and conjugated (*right*). Percent relative abundance was calculated based on area under the peak/total ion current × 100.

For unconjugated bile acids, no major differences were observed between the four solvent types in the pre-extraction buffers (**Figure 2A-2C, left**). Across the second-step dilution conditions, most unconjugated bile acids showed only modest changes in signal intensity, with slightly lower signal in Conditions 2 and 3 compared to Condition 1. This was also true when using the larger volume (100 μL vs. 50 μL) of pre-extraction supernatant. One exception was lithocholic acid (LCA), which exhibited slightly higher intensity under Conditions 2 and 3. Since LCA typically elutes later in our LC conditions compared to many other unconjugated bile acids, it is less affected by co-elution and may be more aided by the reduced matrix interference. This may also explain the higher signal when using less sample material in Conditions 2 and 3. In contrast, the conjugated bile acids showed greater changes with extraction conditions (**Figure 2A-2C, right**). For taurine-conjugated bile acids, choice of pre-extraction buffer led to measurable differences in extraction efficiency. Specifically, using a 1:1 methanol:acetonitrile buffer in the pre-extraction step resulted in slightly higher intensity across the three dilution conditions. Furthermore, conjugated bile acid losses were observed in both Conditions 2 and 3, particularly when methanol or ethanol are used as pre-extraction buffers. Using the matching organic solvent in Condition 3 also caused more signal reduction than Condition 2 (**Figure 2B-2C**). For the glycine-conjugated bile acids, the extraction efficiency followed a similar trend as observed for taurine-conjugated bile acids (**Figure S1**). Thus, among the stool extraction conditions tested, 1:1 acetonitrile/methanol pre-extraction followed by the 1:1 methanol dilution in Condition 1 provided the most consistent overall profile, balancing recovery with detection coverage (**Figure 2A**).

### Serum and Plasma Bile Acid Extraction

Since the stool matrix is particulate-rich and highly variable, we performed a broad comparison of solvent compositions and second-step dilution schemes to identify conditions that preserve both unconjugated and conjugated bile acids. In contrast, serum and plasma are protein-rich, and bile acids are typically quantified from the post-precipitation supernatant following organic-solvent protein precipitation. Thus, rather than optimize the 1:1 acetonitrile/methanol solvent, which is widely used for serum/plasma protein precipitation, we focused on sample input volume as a practical constraint that often limits study designs. We also wanted to include both human serum and plasma because they can differ in matrix composition (e.g., clotting factors and anticoagulants), which may impact recovery.

To extract the bile acids from the human control serum and plasma, the 1:1 acetonitrile/methanol extraction solvent was fixed at 1:4 solvent:sample ratio (**Figure 1**) For example, both 50 μL and 200 μL starting volumes of serum and plasma were utilized to understand recovery differences. This resulted in either 200 μL of solvent added to the 50 μL samples or 800 μL of solvent added to the 200 μL samples. Across both serum and plasma, the higher input volume (200 μL) increased overall signal and improved detectability of low-abundance conjugated bile acids (**Figure 3A-3B, right**). Reducing the sample volume to 50 μL had minimal impact on unconjugated bile acids (**Figure 3A-3B, left**), but reduced detection of some conjugated bile acids. For example, tauroursodeoxycholic acid (TUDCA) became undetectable in serum at 50 μL under the same 1:4 ratio (**Figure 3B**). These results suggest that smaller input volumes can still support broad bile acid profiling but may reduce detection and annotation confidence for low-abundance conjugated bile acids. For glycine-conjugated bile acids, the extraction efficiency followed a similar trend to the taurine-conjugated bile acids, reflecting their comparable extraction behavior (**Figure S2**).

**Figure 3.**
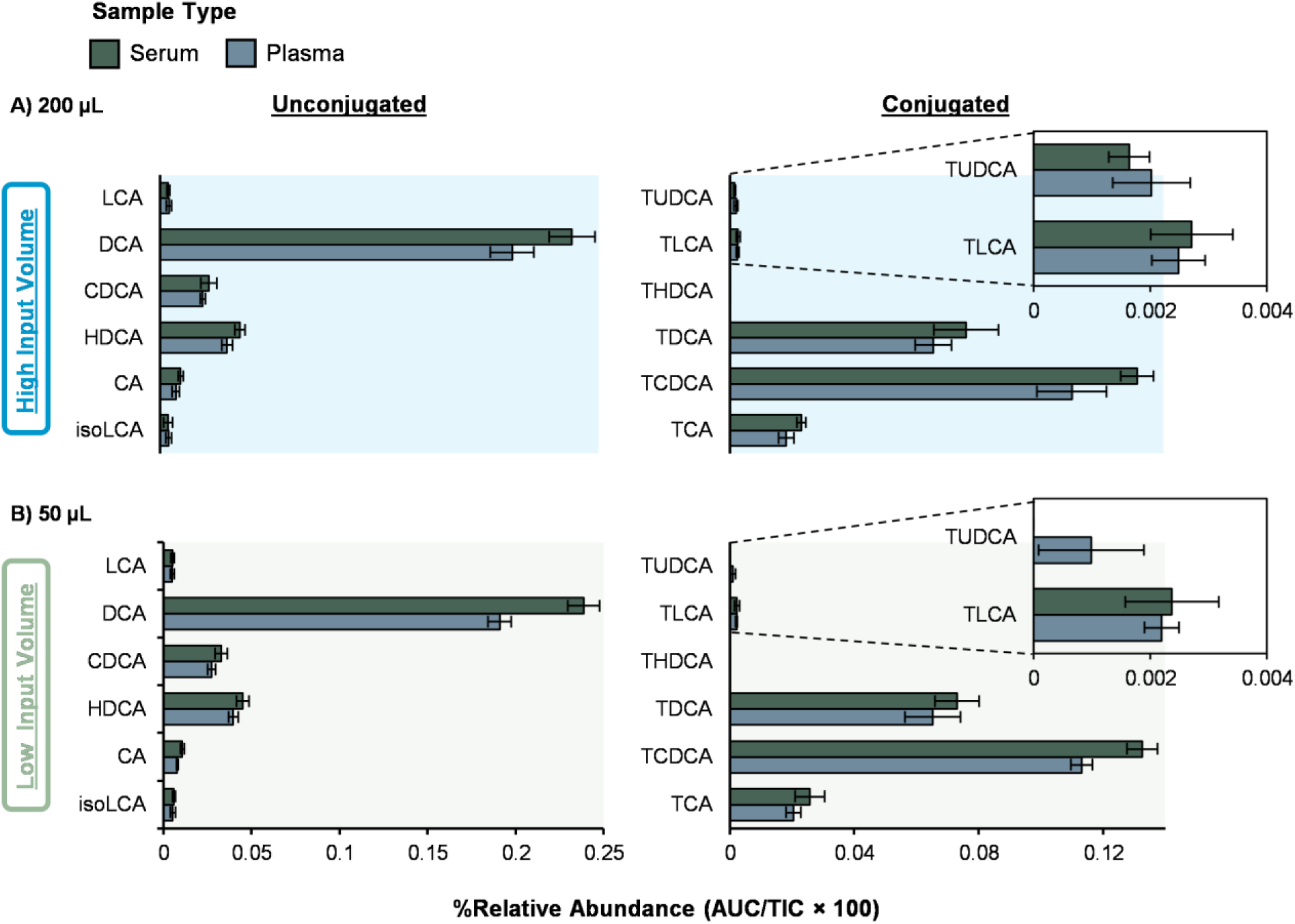
Extraction conditions for human serum and plasma samples. Bile acid extraction performance was evaluated at a fixed 1:4 solvent-to-sample ratio for both **A)** 200 μL and **B)** 50 μL serum and plasma volumes using 1:1 acetonitrile/methanol. A Skyline search against the full library detected 34 bile acids, and a representative subset of 12 are shown with unconjugated (*left*) and conjugated (*right*). Percent relative abundance was calculated based on area under the peak/total ion current × 100.

NIST reference serum and plasma were also processed alongside the study samples for all extraction conditions. In all cases, they showed similar results to the control human samples (**Figure S3**). We also tested an exploratory higher-dilution condition (1:8 with 50 μL) to probe the effect of increased dilution at low sample input (**Figure S4**). The 1:8 condition suggested further loss of taurine-conjugated bile acids and is not recommended. Thus, we recommend the 1:4 extraction using 1:1 acetonitrile/methanol and 200 μL input when available. When sample is limited, 50 μL at the same 1:4 ratio is a practical alternative but may reduce detection and annotation confidence for lower abundance conjugated bile acids.

### LC Separations

Chromatographic separation is critical for confident bile acid annotation as mass spectrometry analysis alone often cannot distinguish bile acid isomers with identical precursor *m/z* values and similar fragmentation. Here, we evaluated the effects of mobile phase pH and stationary phase chemistry using two reverse-phase columns and two bile acid mixtures: (i) a Cayman 18-bile acid mixture containing unconjugated bile acids and glycine- and taurine-conjugated species (catalog no. 33505) and (ii) 22 synthetic amino acid-conjugated cholic acids (AA-CA) as representative MCBAs (**Figures 4-5**). The three sets of mobile phase compositions and their gradient programs are provided in the **Supplemental Information** and **Table S4**. Retention times for bile acids detected under the three LC conditions are also provided in **Tables S1-S2**. Because experiments were acquired across different days, absolute signal intensities are run-specific and intended for comparison within figures rather than across figures (**Figures 4-6**).

**Figure 4.**
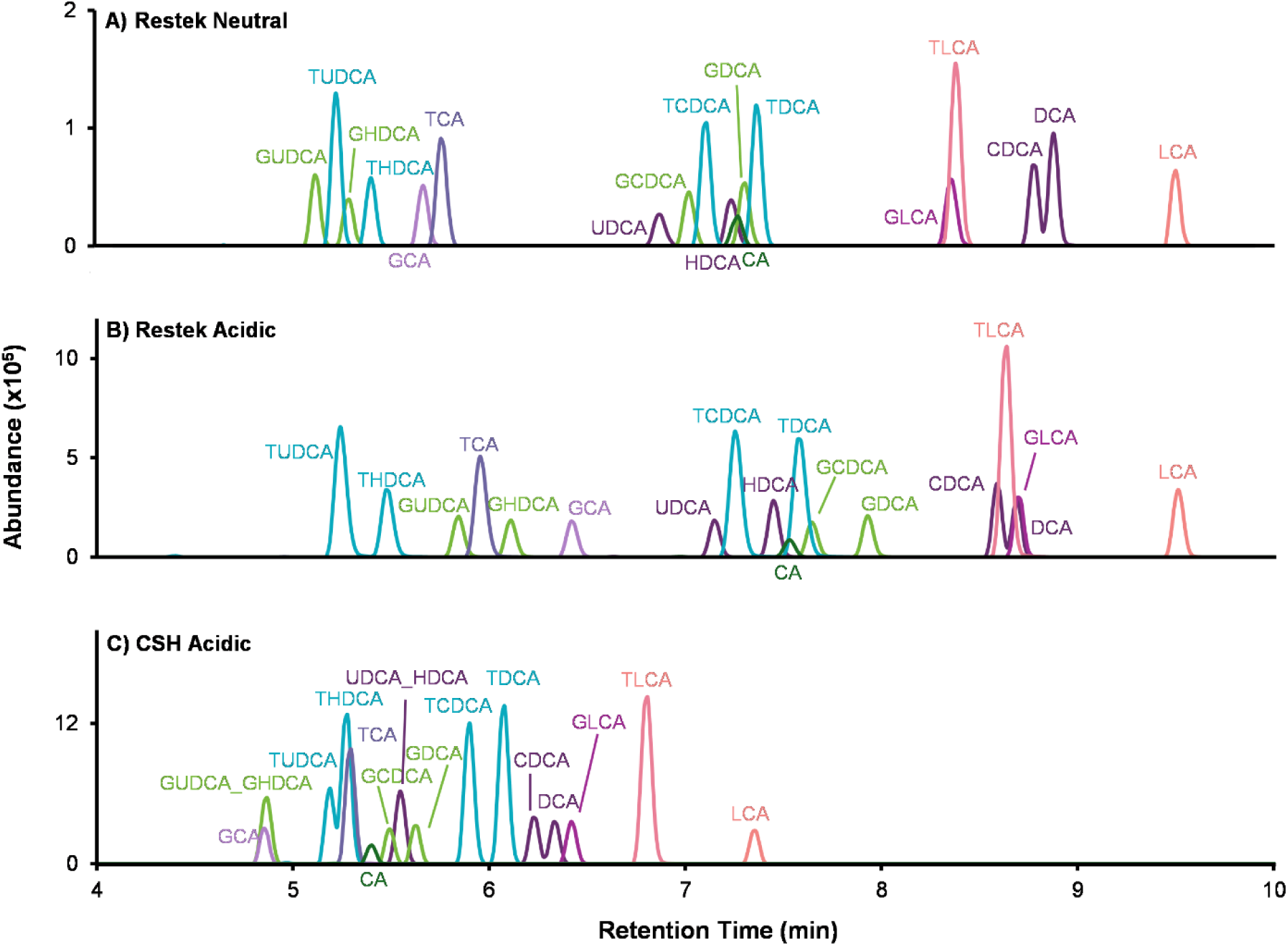
LC effects on bile acid separation. Three LC methods were evaluated using an 18 bile acid standard mixture to assess the effects of mobile-phase pH and stationary phase chemistry. (A) Restek C18 column with a neutral mobile phase. (B) Restek C18 column with an acidic mobile phase containing 0.1% formic acid. (C) CSH C18 column with the acidic mobile phase.

**Figure 5.**
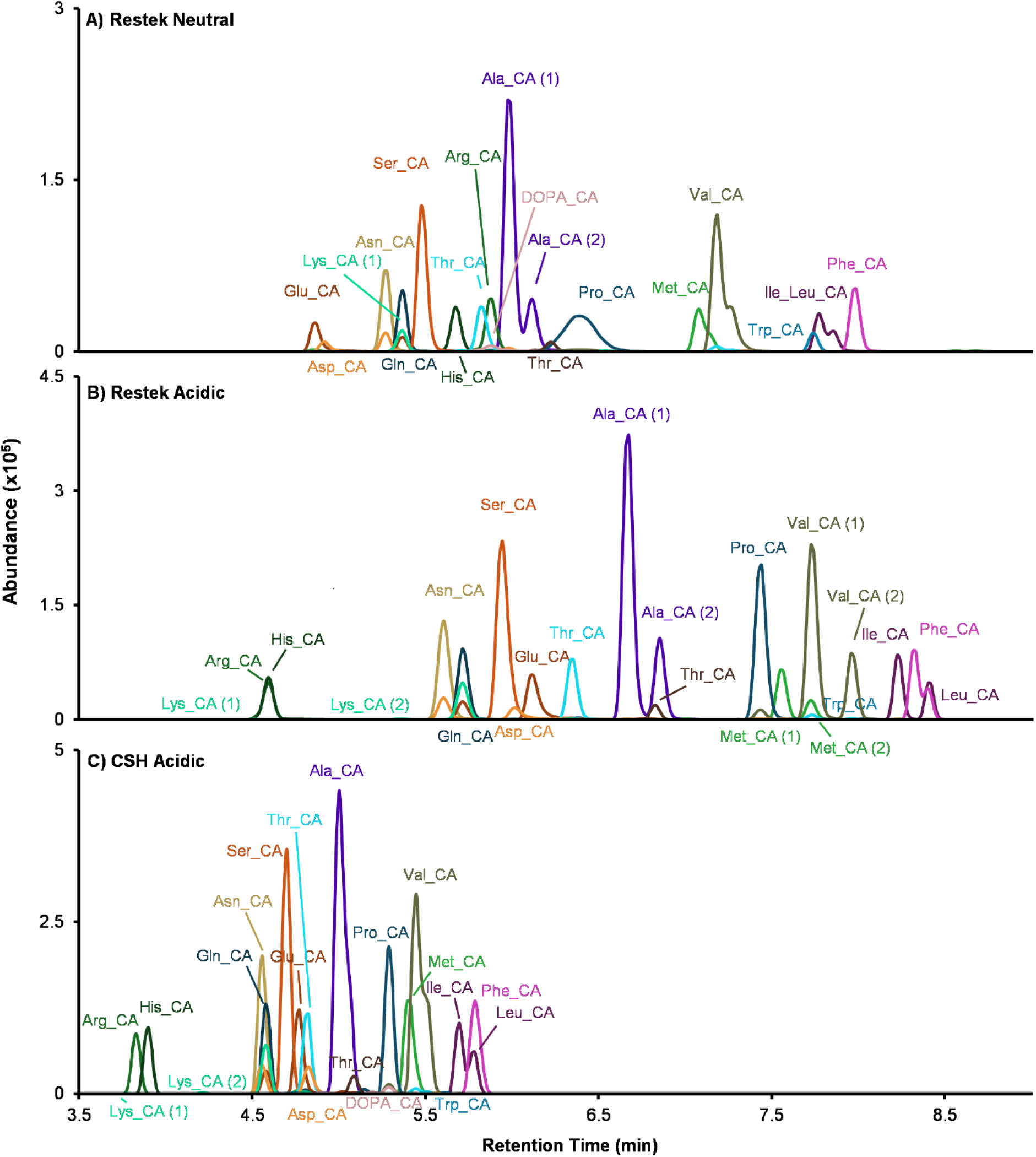
LC effects on separation of amino acid-conjugated cholic acids. Three LC methods were evaluated using 22 amino acid-conjugated cholic acids to compare separation under different mobile-phase pH and stationary phase chemistries. (A) Restek C18, neutral mobile phase. (B) Restek C18, acidic mobile phase containing 0.1% formic acid. (C) CSH C18, acidic mobile phase.

**Figure 6.**
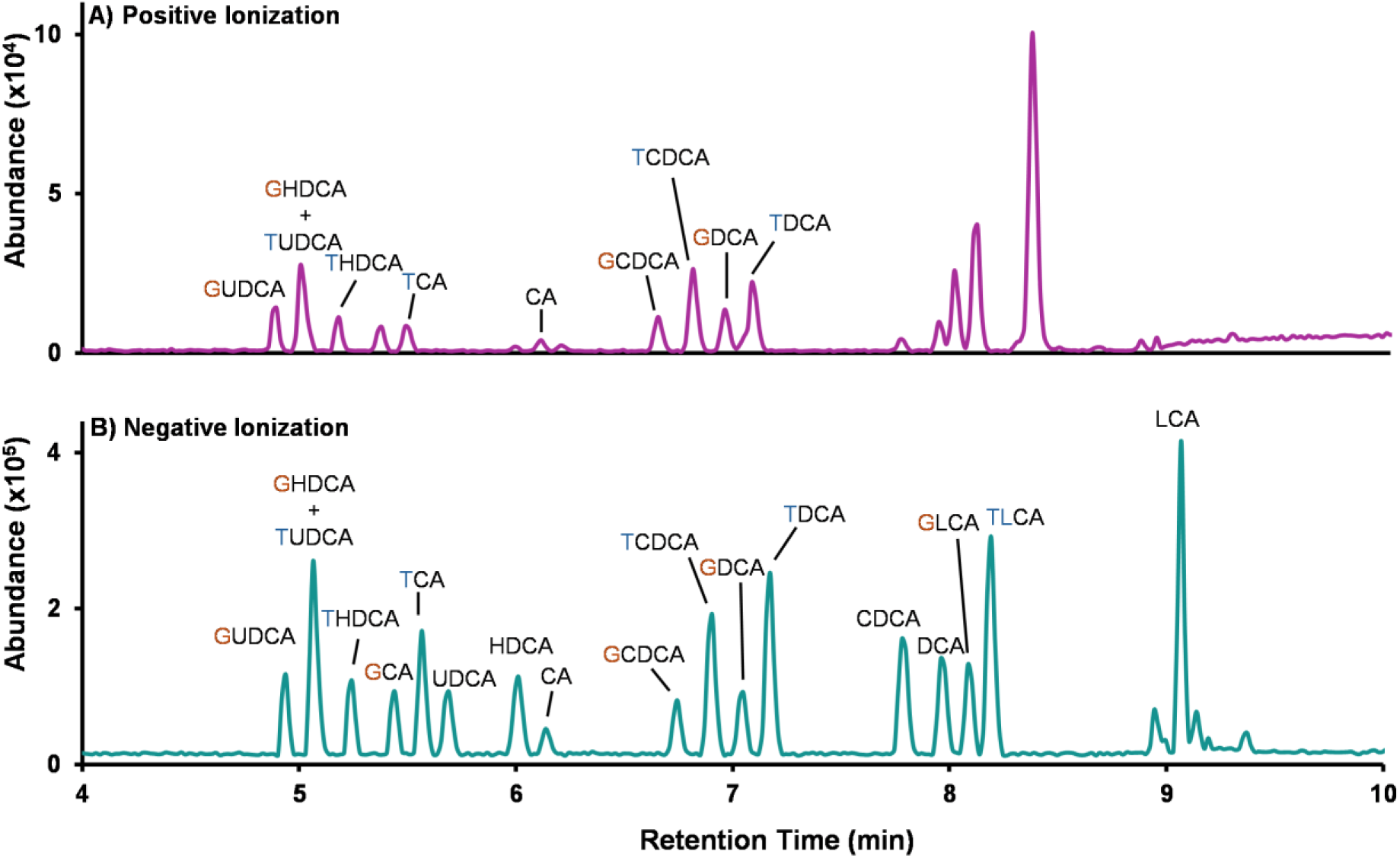
Effect of ionization mode on bile acid detection. Base peak chromatograms acquired in positive (top) and negative (bottom) ionization for the 18 bile acid standard mix. Higher signal intensity and a more uniform response were observed in negative ionization mode. Peaks are labeled by bile acid identity. In positive mode, several bile acids in the 18-component mixture did not produce a detectable signal under the acquisition conditions used. Unlabeled peaks reflect additional signals that do not correspond to the 18 standards, including background ions and other mixture-related features.

We first tested three LC methods using the Cayman standard mix to isolate the effects of mobile phase pH and column chemistry (**Figure 4**). Briefly, as noted in the Methods section, Method 1 used a Restek C18 column with a neutral mobile phase (**Figure 4A**); Method 2 used the same Restek column with an acidic mobile phase, adding 0.1% formic acid to assess pH effects on retention and selectivity (**Figure 4B**); and Method 3 used the same acidic mobile phase on a Waters ACQUITY Premier charged surface hybrid (CSH) C18 column to evaluate whether additional charge interactions on the stationary phase under acidic conditions improve separation of bile acids (**Figure 4C**). This design isolates the effects of mobile phase pH (**Figure 4A** vs. **Figure 4B**) and stationary phase chemistry under acidic conditions (**Figure 4B** vs. **Figure 4C**). On the Restek column, the acidic mobile phase reduced peak overlap compared to the neutral mobile phase, consistent with pH-dependent changes in retention and selectivity on the same stationary phase. By comparison, the CSH column with an acidic mobile phase reduced co-elution for some bile acids (*e.g.,* GLCA with TLCA (*Restek neutral*), and GLCA with DCA (*Restek acidic*)), but reduced separation for the isomer pair UDCA and HDCA and their glycine and taurine conjugated forms. Thus, each column has benefits and limitations.

Since MCBAs include diverse conjugation chemistries, we expected their chromatographic behavior to be sensitive to mobile phase pH and column chemistry. We therefore analyzed a set of 22 AA-CA species across the same three LC methods (**Figure 5**), comparing peak overlap and qualitative separation patterns across methods and monitoring column performance during repeated injections. Across methods, mobile phase pH altered AA-CA elution order and selectivity.

For workflows that require neutral mobile phase conditions for method compatibility or to reduce column stress under extended use, the Restek C18 column with a neutral mobile phase (**Figure 5A**) separated many bile acids. The Restek C18 column with an acidic mobile phase (**Figure 5B**), however, improved separation (*e.g.,* isoleucine- and leucine-conjugated cholic acid (Ile/Leu-CA)), supporting more detailed structural analysis of the bile acids, but showed reduced robustness over extended use as indicated by increasing backpressure and retention time drift. The CSH C18 column with an acidic mobile phase (**Figure 5C**) produced shorter elution times for the amino acid-conjugated cholic acids, with AA-CA eluting before 6 min, reducing overlap with the unconjugated bile acids relative (**Figure 5C** vs **Figure 4C**), Additionally, the shorter LC gradient time supports this method as an efficient option for higher-throughput analyses.

Across the 18 bile acid standard mixture, acidic conditions generally increased detection signal under our tested methods, whereas AA-CA signals were broadly similar across neutral and acidic conditions. Because chromatographic selectivity and elution order also changed between conditions, we describe these differences as method-dependent rather than attributing them solely to intrinsic ionization efficiency. Therefore, selection of mobile phase and column should balance isomer separation and long-term robustness and be tailored to specific study objectives and throughput requirements. Thus, our observations are that the Restek C18 column with acidic mobile phase conditions provided improved separation for several targets but placed greater stress on the column under our operating condition, which can affect long-term stability, as evidenced by backpressure increases and retention time drift across repeated injections. For studies requiring neutral pH conditions or prioritizing long-term robustness, the Restek C18 column with a neutral mobile phase may be the preferred choice, despite its lower separation performance. For large-scale studies where maximizing chromatographic separation and reducing peak co-elution is not the primary objective, the CSH C18 column under acidic mobile phase offers a higher throughput option and maintained more stable backpressure, indicating better tolerance to acidic operation.

### Ionization Mode Selection

Ionization polarity strongly influences bile acid detectability and annotation (**Figure 6**). We compared positive and negative polarities with ESI using the Cayman Standard Mixture containing 18 bile acids which include unconjugated and taurine- and glycine-conjugated species (catalog no. 33505). These comparisons were performed using Method 1 on the Restek C18 column with a neutral mobile phase to provide a consistent baseline condition for evaluating polarity-dependent ion formation. Under the tested conditions, negative ion mode exhibited higher signal intensities (approximately 8-fold greater) across most tested bile acids, consistent with improved detectability in negative mode. Moreover, negative mode provided a more uniform ionization response and cleaner background in the base peak chromatogram across bile acid species, which is advantageous for profiling workflows that aim to capture both high-and low-abundance species.

In positive mode, bile acids formed protonated, sodiated, and ammoniated ions ([M+H]⁺, [M+Na]⁺, and [M+NH₄]⁺), which can provide complementary evidence for annotation when matched to the reference library. However, because adduct distributions vary by structure and acquisition conditions, signal can be distributed across multiple ion forms, which complicates consistent quantification when different adducts dominate for different bile acids. This resulted in not all bile acids in the standard mixture producing a detectable signal and additional non-target features being observed in the base peak chromatogram (**Figure 6A**). Therefore, we prioritized negative mode for primary profiling and quantification, and we treat positive mode as a complementary polarity for additional confirmation when needed. **Tables S1-S2** list features detected in each polarity and their observed ion forms, enabling users to determine whether a target bile acid is detected for a given mode and whether multiple adducts can be used to support annotation.

### Multidimensional Bile Acid Library

To support bile acid annotation and structural characterization, we created a multidimensional LC-IMS-MS reference library containing LC retention times, experimental drift time IMS CCS values in nitrogen (^DT^CCS_N2_), and accurate masses for 280 unique bile acids with each ion form and adduct annotation detected (**Tables S1-S2**). Chemical structure information and identifiers for each bile acid are also included in **Table S3**. In total, 18.9 % (53 of 280 unique bile acids) were structures that were not yet present in PubChem (new CIDs 177860373-177860425 were created upon deposition of the structures). The library comprises 343 deprotonated features in negative ionization mode and 665 features (including sodiated, ammoniated and protonated species) in positive mode. When multiple features corresponded to the same bile acid identity, we assigned suffixes based on separation dimension: where (1) or (2) indicate LC-resolved features separated by retention time, whereas (a) or (b) indicates features separated primarily by IMS. This convention is used consistently throughout **Tables S1-S2**. To illustrate trends observed in the bile acids, CCS versus *m/z* relationships for the library are shown for negative mode (**Figure 7A**) and positive mode (**Figure 7B**, and **Figure S5** for different ion types). Across ionization modes and adduct types, bile acids show distinct groupings by conjugate class. For example, glycine- and taurine-conjugated bile acids form separable clusters in CCS versus *m/z* space. Thus, this multidimensional library should improve discrimination of structurally similar bile acids, including isomers and amino acid-conjugated species, in complex biological samples.

**Figure 7.**
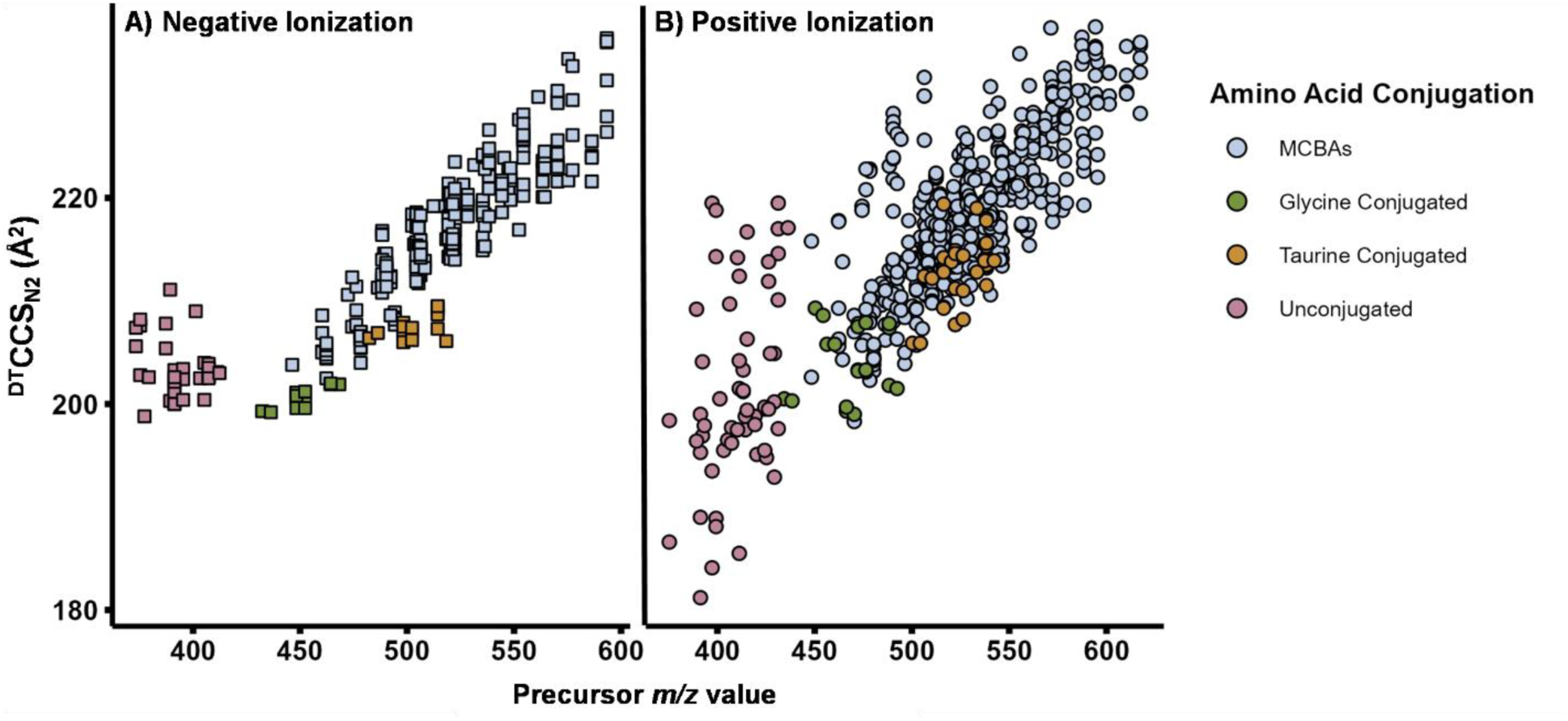
Bile acid library collision cross section (CCS) values versus mass-to-charge ratio (*m/z*) trends across ionization modes. A) CCS values were measured for bile acids under negative ionization mode resulting in [M-H]^-^ ions and **B)** positive ionization mode resulting in [M+H]^+^, [M+Na]^+^, and [M+NH_4_]^+^ ions. The bile acids are grouped by unconjugated, glycine and taurine conjugates, and MCBAs in the plots to illustrate the different trends.

## Conclusion

In this work, we evaluated key analytical variables that influence bile acid detectability and annotation across complex biological matrices including stool, serum and plasma. Specifically, we compared extraction workflows, assessed chromatographic conditions and their practical trade-offs in separation performance and robustness, and evaluated ionization behavior across polarities. Across extraction conditions, unconjugated bile acids were comparatively stable, whereas conjugated bile acids were more sensitive to workflow choices, underscoring the importance of solvent selection and handling steps when low-abundance conjugated species are of interest. For serum or plasma, we held the precipitation solvent system constant and focused on practical constraints such as available sample volume, showing that higher input improved detectability of lower abundance conjugated bile acids under the tested conditions. Three LC methods were also evaluated for the bile acids with each showing advantages and limitations. Collectively, these comparisons highlight that analytical conditions depend on the matrix and study goals, and that documenting solvent composition, dilution strategy, and input amounts is important for reproducible bile acid profiling.

A multidimensional LC-IMS-MS reference library for 280 unique bile acids that integrate structures, LC retention time, accurate mass, experimental CCS values across ionization modes and annotated ion forms and adduct types was also created in this work. The library includes 343 deprotonated features in negative mode and 665 features in positive mode spanning common adduct forms (e.g., protonated, sodiated and ammoniated). By providing orthogonal separation information alongside curated feature expectations, this resource improves discrimination of structurally similar bile acids, including isomers and microbially conjugated species, and supports more confident annotation in complex samples. Together, the evaluated workflows and accompanying reference library provide a foundation for more consistent bile acid profiling and support future studies of bile acid diversity across host and microbiome contexts.

## Sustainability Statement

This article aligns with UN SDG3 (One Health: Good health and wellbeing) by enabling more comprehensive and accurate analytical measurements of bile acids, supporting deeper investigations into their biological roles and disease associations. These improved evaluations will also inform the development of better diagnostics and therapeutics to advance human health.

## Acknowledgements

Funding for this work was made possible through grants from the NIH National Institute of Environmental Health Sciences (P42 ES027704) and the National Institute of General Medical Sciences (R01 GM141277 and RM1 GM145416). This research was also supported [in part] by the National Center for Biotechnology Information of the National Library of Medicine (NLM), National Institutes of Health (NIH). The contributions of the NIH author(s) are considered Works of the United States Government. The findings and conclusions presented in this paper are those of the author(s) and do not necessarily reflect the views of the NIH or the U.S. Department of Health and Human Services.

