## Supplementary_Information for "Unlocking the Bile Acid Universe: Advanced Workflows and a Multidimensional Library of 280 Unique Species"

Table of Contents

Cover page and table of contents	**S1-2**

**Supplemental Methods**	**S3**

UCSD Standard Preparation and Subsequent LC-IMS-MS Analysis	**S3**

Bile Acid Extraction from Solid Matrices	**S4-6**

Bile Acid Extraction from Liquid Matrices	**S7-9**

LC Conditions and Recommended Run Order	**S10**

Optional Quantification Guidance	**S11-12**

**Figure S1.** Stool extraction for glycine-conjugated bile acids	**S13**

**Figure S2.** Serum and plasma extraction for glycine-conjugated bile acids	**S14**

**Figure S3.** Serum and plasma extraction for bile acids under 1:4 (v/v) extraction conditions	**S15**

**Figure S4.** Higher-dilution extraction condition 1:8 (v/v) for bile acids	**S16**

**Figure S5.** Representative collision cross section (CCS) distributions for bile acid features detected in positive-ion mode	**S16**

**Supplemental Methods**

**UCSD Standard Preparation and Subsequent LC-IMS-MS Analysis**

***UCSD Standard Preparation.*** Additional standards were obtained from the Dorrestein lab (UCSD) and analyzed to expand the LC-IMS-MS reference library. All UCSD standards were supplied as powders. Upon arrival, each powdered standard was reconstituted in 1000 μL methanol to prepare an individual stock solution. Stocks were vortex-mixed until fully dissolved and stored for subsequent analyses. Details of the UCSD standards are provided in Table S9.

***Flow Injection Analysis (FIA).*** Stock standard solutions were diluted in 50:50 (v/v) methanol:nanopure water for MS analysis. FIA (direct injection) was used to rapidly evaluate ionization behavior and IMS features for each UCSD standard in both ionization modes. For FIA, 0.01-0.2 μL of standard solution was injected in triplicate using a 50:50 mixture of mobile phase A (water + 5 mM ammonium acetate) and mobile phase B (50:50 (v/v) methanol:acetonitrile) (mobile phase set 1). Data were acquired in both positive and negative ionization modes over *m/z* 50-1700.

***LC-IMS-MS Analysis.*** UCSD standards were subsequently analyzed by LC-IMS-MS under the chromatographic methods described in the main Methods. In negative ionization mode, each standard was injected once using the Restek neutral method and once using the CSH acid method. In positive ionization mode, each standard was injected once using the Restek neutral method.

***Data Processing.*** Raw .d files were demultiplexed using PNNL-Preprocessor 4.0 with a moving average of 3, a minimum pulse coverage of 100%, and a signal intensity threshold of 20 counts, generating DeMP.d files. Demultiplexed data were analyzed in Agilent MassHunter IM-MS Browser 10.0 to extract drift times. CCS values were then calculated using the single-field approach in a Microsoft Excel workbook, which applies the single-field calibration parameters to convert drift time to CCS. Microsoft Excel was also used to summarize mass accuracy (ppm error of measured *m/z*) and CCS precision across replicate injections. For the UCSD standards reported here, m/z errors were under 10 ppm and CCS %RSD across replicates was below 0.3%.

**Bile Acid Extraction SOP**

This Standard Operating Procedure (SOP) provides general guidance for extracting bile acids from solid and liquid biological matrices for downstream analysis by LC-IMS-MS (and conventional LC-MS, where applicable). The workflow is designed to be adaptable across sample types, spanning solid matrices such as feces, cecal contents, liver, and other tissues, and liquid matrices such as serum, plasma, bacterial culture supernatants, and other biofluids. The procedures use matrix-appropriate homogenization or protein-precipitation strategies, internal standardization, and optional external calibration for absolute quantification. Key parameters (e.g., solvent composition, solvent-to-sample ratio, internal standard concentration/spike volume, and clarification/filtration conditions) should be optimized as needed to accommodate matrix or sample dependent properties (e.g., expected bile-acid abundance) and the analytical objectives. Chromatographic conditions and a recommended sample injection sequence are provided to support method implementation and batch quality control.

**Bile Acid Extraction from Solid Biological Matrices including Stool**

**Materials and Reagents**

*1. Make Internal Standard (ISD):*

- Concentration**:** 50-200 μM solution in 50% methanol (MeOH), depending on experimental needs
- Catalog Numbers**:** MSK-BA1 and MSK-BA2 from Cambridge Isotope Laboratories
- Preparation (as the example of 200 μM final concentration)**:**
  i. Add 250 μL of 50:50 (v/v) methanol:nanopure water to each ISD vial.
  ii. Vortex until the contents are fully dissolved.
  iii. Centrifuge the vials at 500 x g for 1 minute.
  iv. Combine the contents of both vials into a single ambient container from Agilent. (Part number: 8010-0543)

*2. Make Unlabeled Standard:*

- Concentration**:** 100 μM solution in 50% methanol (MeOH)
- Catalog Numbers**:** MSK-BA1-US-1 and MSK-BA2-US-1
- Preparation**:**
  i. Add 500 μL of 50:50 (v/v) methanol:nanopure water water to each vial.
  ii. Vortex until the contents are fully dissolved.
  iii. Centrifuge the vials at 500 x g for 1 minute.
  iv. Combine the contents of both vials into a single ambient vial from Agilent. (Part number: 8010-0543)

***Note:*** A 100 µM stock is recommended to support calibration that spans higher bile acid abundances. If lower concentration standards are needed, they can be prepared by serial dilution from the 100 µM stock using 50:50 (v/v) MeOH:water. Alternatively, a commercial bile acid mixture at lower concentration (e.g., Bile Acids MaxSpec® Discovery Mixture, Cayman Chemical) may be used directly when the expected concentration range does not require high-end coverage.

*3. Prepare an Extraction Solvent: 1:1 acetonitrile (ACN): methanol (MeOH) with phosphate (Pre-extraction buffer)*

- Composition: 1:1 (v/v) acetonitrile (ACN):methanol (MeOH) containing phosphate (final 3 mM)
- Stock Potassium Phosphate Buffer: 100 mM (Catalog No.: 7778-77-0)
- Preparation**:**
  i. Dilute the 100 mM phosphate stock 1:5 (v/v) with water to obtain 20 mM phosphate.

ii. Mix 60 mL of 20 mM phosphate buffer with 340 mL organic solvent (170 mL MeOH + 170 mL ACN) to yield 400 mL of pre-extraction buffer with 3 mM phosphate.

*4. NIST Standard for Reference*

- NIST 1950 Metabolites in human plasma
- NIST 909C Human serum

*5. LC Vials: Agilent glass vials with inserts*

- Catalog Number: 5188-6591 from Agilent

*6. Optima* *™LC-MS Grade Solvents for Mobile Phases and Blanks*

- Methanol: Catalog Number: 67-56-1
- Acetonitrile: Catalog Number: 75-05-8
- Water: Catalog Number: 7732-18-5

**All solvents purchased from Fisher Chemical, Pittsburgh, PA*

*7. Sample Tubes for Extraction*

- 1.5 mL microcentrifuge tube: Catalog Number: 3451 (Thermo Scientific)
- 2 mL beadmill tube: Catalog Number: 19-620 (Revvity)
- PTFE spin filter tube: Catalog Number: UFC30LG25 (Millipore)

**Extraction Procedure for Solid Matrices:**

1. Internal Standard Addition

Add 2 μL of 200 μM ISD solution (#1 in Materials) to each 2 mL beadmill tube (#7 in Materials) containing 50-100 mg solid sample (e.g., stool).

2. Sample-to-solvent Weight Normalization

Add extraction buffer (#3 in Materials) at 8 µL per 1 mg sample (8:1 µL/mg).

Example: 75 mg sample, add 600 µL extraction buffer.

3. Homogenization

Homogenize using a Fisherbrand™ Bead Mill 24 Homogenizer (Hampton, NH, USA; catalog number 15-340-163) at 2.1 m/s for 2 minutes in a single cycle

***Note:*** This setting is typically sufficient for fecal and cecal samples. For harder tissues (e.g., liver or other solid organs), two homogenization cycles may be required. Allow a brief pause, placing tubes on ice between cycles, to minimize sample heating.

4. Centrifugation

Centrifuge the samples at 10,000 × g for 15 min or 13,000 × g for 10 min at 4°C.

***Note:*** For solid matrices, lower centrifugation speeds may be preferable when tube balancing is limited by sample mass variability.

5. Extraction

Transfer 100 µL of supernatant into a fresh 1.5 mL microcentrifuge tube and add 100 µL MeOH. Mix briefly.

6. Mixing

Mix at 600 rpm for 20 min at room temperature using an Eppendorf MixMate® (Catalog No.: 5353000529).

7. Filtration

Transfer the full 200 µL to a PTFE spin filter tube and centrifuge at 10,000 × g for 1 min at 4°C. Collect the filtrate.

8. Preparation for LC-MS Analysis

Transfer 60 µL filtrate into labeled LC vials (#5 in Materials) for LC–IMS–MS analysis (or LC–MS, where applicable).

***Note:*** If needed to improve solvent compatibility with the initial LC conditions, the extract may be diluted with nanopure water. In our workflow, extracts were injected without dilution and acceptable chromatography was obtained; dilution may be used as an optimization step if peak shape or retention is suboptimal. Record any dilution factor applied, as this decreases analyte concentration.

**Bile Acid Extraction from Liquid Biological Matrices including Serum and Plasma**

**Materials and Reagents**

*1. Make Internal Standard (ISD):*

- Concentration**:** 50-200 μM solution in 50% methanol (MeOH), depending on experimental needs
- Catalog Numbers**:** MSK-BA1 and MSK-BA2 from Cambridge Isotope Laboratories
- Preparation (as the example of 200 μM final concentration)**:**
  i. Add 250 μL of 50:50 (v/v) MeOH:watermethanol:nanopure water to each ISD vial.
  ii. Vortex until the contents are fully dissolved.
  iii. Centrifuge the vials at 500 x g for 1 minute.
  iv. Combine the contents of both vials into a single ambient container from Agilent. (Part number: 8010-0543)

*2. Make Unlabeled Standard:*

- Concentration**:** 100 μM solution in 50% methanol (MeOH)
- Catalog Numbers**:** MSK-BA1-US-1 and MSK-BA2-US-1
- Preparation**:**
  i. Add 500 μL of 50:50 (v/v) MeOH:watermethanol:nanopure water water to each vial.
  ii. Vortex until the contents are fully dissolved.
  iii. Centrifuge the vials at 500 x g for 1 minute.
  iv. Combine the contents of both vials into a single ambient vial from Agilent. (Part number: 8010-0543)

***Note:*** A 100 µM stock is recommended to support calibration that spans higher bile acid abundances. If lower concentration standards are needed, they can be prepared by serial dilution from the 100 µM stock using 50:50 (v/v) MeOH:water. Alternatively, a commercial bile acid mixture at lower concentration (e.g., Bile Acids MaxSpec® Discovery Mixture, Cayman Chemical) may be used directly when the expected concentration range does not require high-end coverage.

*3. Prepare an Extraction Solvent: 1:1 acetonitrile (ACN): methanol (MeOH)*

- Composition: 1:1 (v/v) acetonitrile (ACN):methanol (MeOH)

*4. NIST Standard for Reference*

- NIST 1950 Metabolites in human plasma
- NIST 909C Human serum

*5. LC Vials: Agilent glass vials with inserts*

- Catalog Number: 5188-6591 from Agilent

*6. Optima* *™LC-MS Grade Solvents for Mobile Phases and Blanks*

- Methanol: Catalog Number: 67-56-1
- Acetonitrile: Catalog Number: 75-05-8
- Water: Catalog Number: 7732-18-5

**All solvents purchased from Fisher Chemical, Pittsburgh, PA*

*7. Sample Tubes for Extraction*

- 1.5 mL microcentrifuge tube: Catalog Number: 3451 (Thermo Scientific)

**Extraction Procedure for Liquid Matrices:**

1. Internal Standard Addition

Add 2 μL of 200 μM ISD solution (#1 in Materials) to each 1.5 mL microcentrifuge sample tube(#7 in Materials).

2. Sample Preparation

Put 50-100 μL of liquid sample (e.g., plasma/serum) to the tube.

3. Solvent Addition

Add 400 µL of extraction solvent (1:1 ACN:MeOH; #3 in Materials) to achieve a 4:1 solvent-to-sample ratio (v/v).

4. Mixing and Sonication

i. Mix at 2,000 rpm for 5 min using an Eppendorf MixMate® (Catalog No.: 5353000529) at room temperature.

ii. Sonicate for 15 min in an ice bath to enhance extraction.

5. Cold Precipitation

Store the samples at -20°C overnight to promote protein precipitation and improve supernatant clarity.

***Note:*** This step may be shortened (e.g., ≥1h at -20°C) depending on matrix protein content and throughput requirements.

6. Centrifugation

Centrifuge at 13,000 × g for 10 min at 4°C.

7. Supernatant Collection

Transfer 300 μL of the supernatant to a new 1.5 mL microcentrifuge tube (#7 in Materials).

8. Preparation for LC-MS Analysis

Transfer 60 µL of the supernatant into LC vials (#5 in Materials) for LC–IMS–MS analysis (or LC–MS, where applicable).

***Note:*** If needed to improve solvent compatibility with the initial LC conditions, the extract may be diluted with nanopure water. In our workflow, extracts were injected without dilution and acceptable chromatography was obtained; dilution may be used as an optimization step if peak shape or retention is suboptimal. Record any dilution factor applied, as this decreases analyte concentration.

**LC Conditions and Recommended Run Order:**

**Mobile Phases**

- Method 1:
- Mobile Phase A1: Water + 5 mM ammonium acetate
- Mobile Phase B1: 50:50 (v/v) methanol:acetonitrile
- Method 2:
- Mobile Phase A2: Water + 0.1% (v/v) formic acid
- Mobile Phase B2: 50:50 (v/v) methanol:acetonitrile + 0.1% (v/v) formic acid
- Method 3:
- Mobile Phase A3: Water + 0.1% (v/v) formic acid
- Mobile Phase B3: Acetonitrile + 0.1% (v/v) formic acid

**LC Columns and Operating Conditions**

- Methods 1 and 2:
  - Restek Raptor C18 (1.8 μm, 50 × 2.1 mm, Catalog No. 9304252)
  - Flow rate: 0.5 mL/min
  - Column temperature: 60°C
  - Mobile phases for Methods 1 and 2
- Method 3:
  - Waters ACQUITY Premier CSH C18 (1.7 μm, 2.1 × 100 mm, Catalog No. 186009461)
  - Flow rate: 0.4 mL/min
  - Column temperature: 60°C
  - Mobile phase: set3

**LC Gradients**

- Detailed gradients are provided in Table S4

**Recommend Sample Injection Sequence (Worklist):**

| **Order** | **Sample** | **Note** |
| --- | --- | --- |
| 1 | Tune Mix | Agilent Tuning Mix (p/n G1969-85000) |
| 2 | Non-injection | Run the LC gradient without injection to equilibrate the system and assess baseline |
| 3 | ES/DB | Injection of the extraction solvent used for sample preparation |
| 4 | EB | Extraction blank, processed using the extraction workflow without biological sample |
| 5 | MB | Method blank, LC-MS grade water processed identically to samples |
| 6 | Unlabeled BA | Unlabeled standard mix (#2 in Materials), injected at an appropriate concentration for system suitability |
| 7 | Samples | Randomized study samples; include NIST reference material and/or calibration standards if absolute quantification is performed |

**Optional Quantification Guidance**

The following guidance is provided as an optional framework for users who wish to implement relative or absolute quantification. Quantification procedures were not a focus of the current study and are not required to reproduce the extraction workflow described here.

*Relative Quantification Using Internal Standards or Absolute Quantification Using External Calibration*

- **Internal standards (recommended for all analysis):**

Stable isotope labeled internal standards (ISD) are spiked into each sample prior to extraction to normalize instrument response, monitor extraction performance, and assess matrix effects across the batch. This approach supports relative comparisons of bile acid abundances between samples.

- **External calibration (for absolute quantification):**

For absolute quantification, prepare an external calibration curve using the unlabeled standard and spike with a fixed volume of the internal standard (ISD). Calibration curves are generated by plotting the peak area ratio of each unlabeled bile acid to the matched deuterated internal standard versus the nominal concentration of the native analyte, and sample concentrations are calculated using the same area-ratio approach. Calibration standards should be processed using the same extraction and cleanup workflow as study samples to minimize bias from matrix effects and sample preparation.

*Steps for Preparation and Processing Calibration Curve Standards*

1. Recommended Standard Concentrations (unlabeled, #2 in Materials):
   - Prepare calibration standards at the following concentrations: 0, 0.16, 0.8, 4, 20, 50, 100 μM.

***Note:*** Concentration points may be adjusted, provided the range spans the expected sample concentrations.

2. Preparation of Calibration Standards:
   - Aliquot 100 µL of each calibration standard into a 1.5 mL microcentrifuge tube.
   - Spike each tube with a fixed volume of internal standard (ISD; #1 in Materials), typically 2 µL (range 2-5 µL depending on ISD concentration and expected instrument response).
   - Mix thoroughly.
3. Processing of Calibration Standards:
   - Process each calibration standard using the same extraction, mixing, and filtration steps as the study samples to ensure equivalent matrix handling and to minimize bias from sample preparation.
   - Calibrators should be randomized within the batch and injected periodically as needed to monitor drift.
   - Matrix-matched calibrators are recommended when feasible to account for matrix effects.

*Extraction Blank and Method Blank*

Process all the blanks using the same steps as experimental samples to ensure comparable handling.

- **Extraction Blank:**

Follow the extraction procedure using an empty extraction tube (no biological sample). Add extraction buffer according to the procedure, but do not add internal standard.

- **Method Blank:**

Follow the Stool Sample Extraction Procedure, Follow the extraction procedure, substituting LC-MS grade water (#6 in Materials) for the biological sample. Add internal standard and extraction buffer as specified in the procedure.

**Figure S1.** Stool extraction for glycine-conjugated bile acids. Bile acid extraction was performed on stool samples using various pre-extraction buffers and second-step dilution conditions. Extraction conditions included four pre-extraction buffers (ethanol, methanol, acetonitrile, or 1:1 acetonitrile/methanol), followed by three dilution conditions: **A)** 1:1 dilution in methanol, **B)** 1:4 dilution in methanol, and **C)** 1:4 dilution in the matching pre-extraction solvent. Glycine-conjugated bile acids were analyzed by LC-IMS-MS. Signal intensity was measured for each condition, with results shown in the corresponding chromatograms and peak areas. Percent relative abundance was calculated based on area under the peak ÷ total ion current × 100.

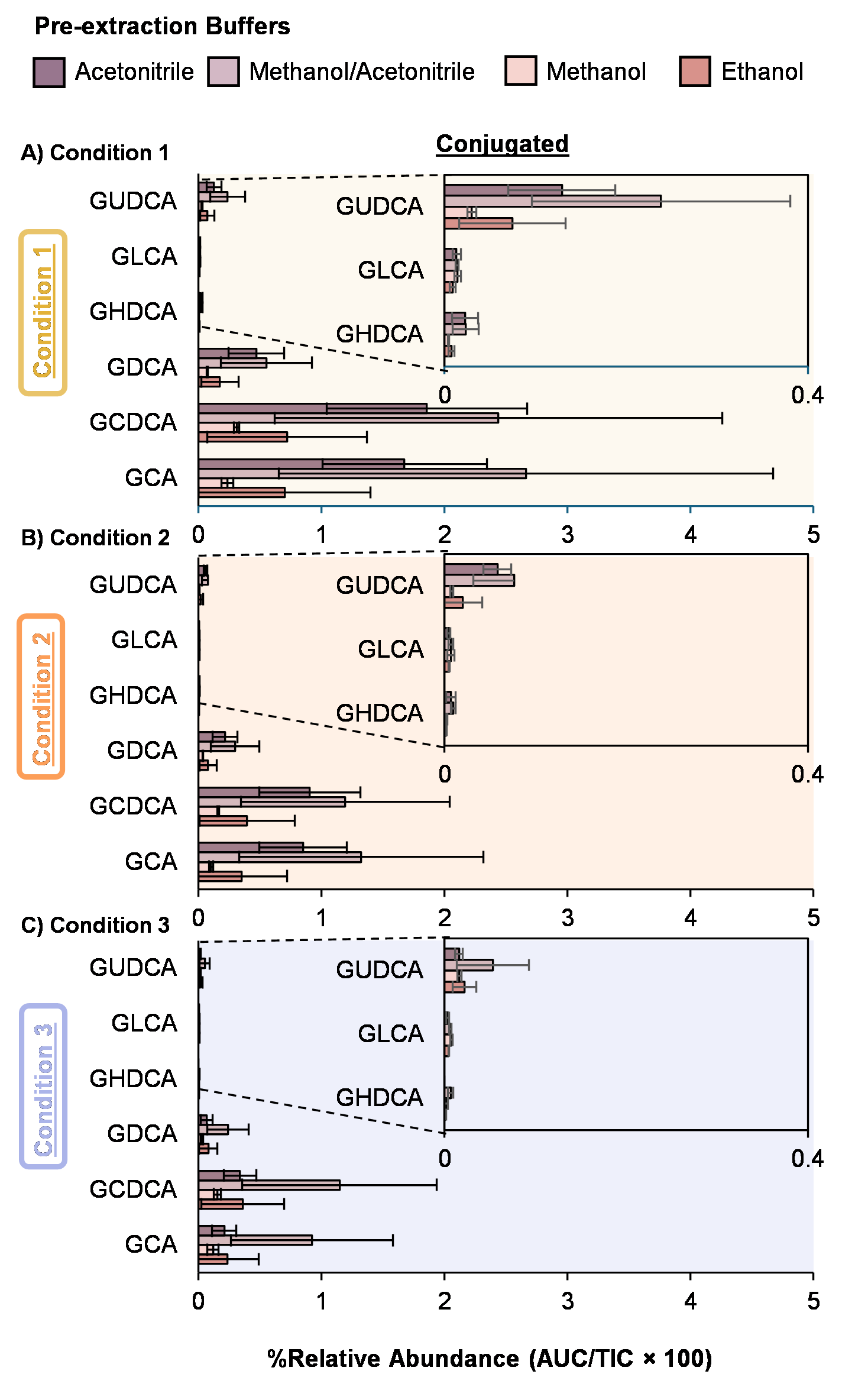

**Figure S2.** Serum and plasma extraction for glycine-conjugated bile acids. Glycine-conjugated bile acids were extracted from human serum and plasma samples using a 1:4 solvent-to-sample ratio with a 1:1 acetonitrile/methanol mixture. Two sample volumes were tested: **A)** 200 μL and **B)** 50 μL. Percent relative abundance was calculated based on area under the peak ÷ total ion current × 100.

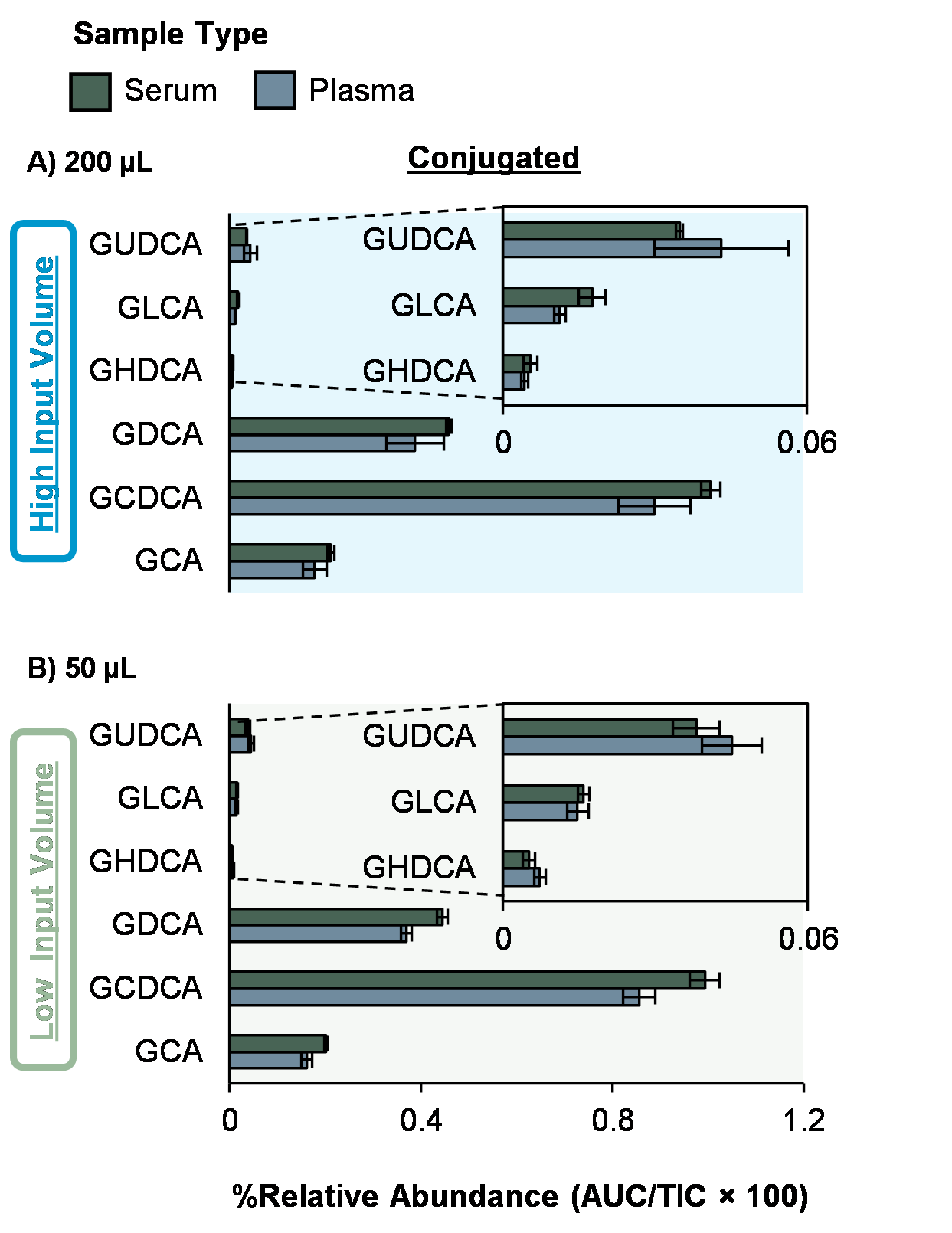

**Figure S3.** Serum and plasma extraction for bile acids under 1:4 (v/v) extraction conditions. NIST reference serum and plasma were extracted using a 1:4 solvent-to-sample ratio with a 1:1 acetonitrile:methanol mixture. Two sample volumes were tested: **A)** 200 μL and **B)** 50 μL. Unconjugated (left), glycine-conjugated (middle), and taurine-conjugated (right) bile acids were analyzed. Percent relative abundance was calculated based on area under the peak ÷ total ion current × 100.

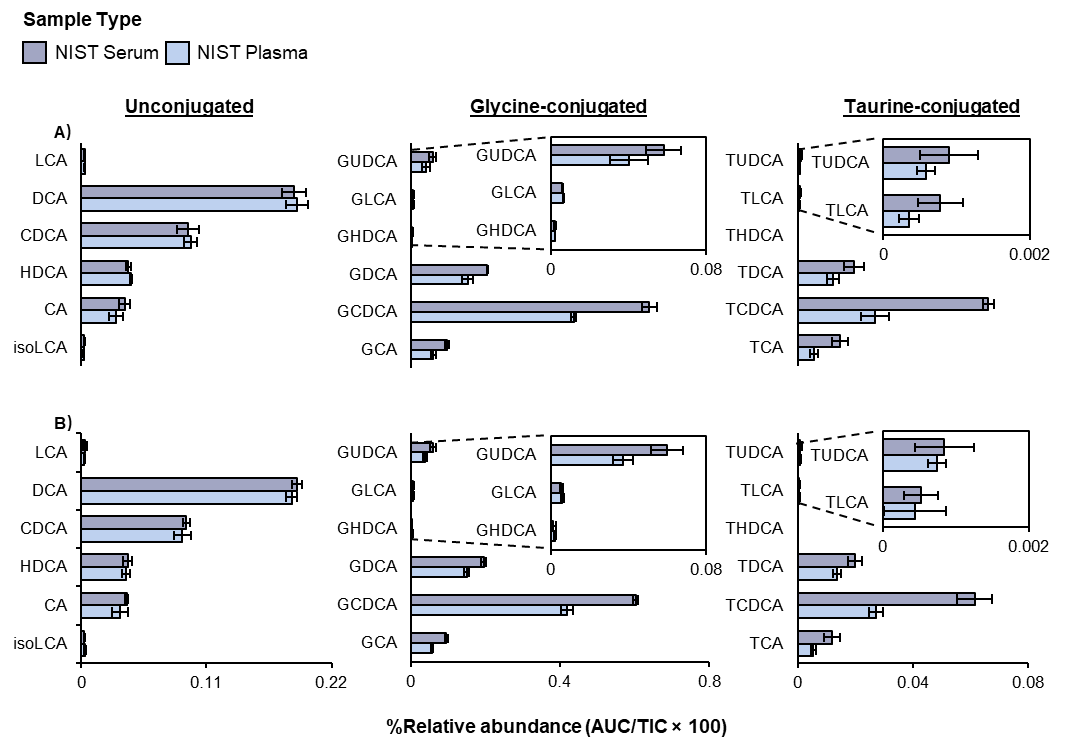

**Figure S4.** Higher-dilution extraction condition 1:8 (v/v) for bile acids. A higher-dilution extraction condition (1:8 solvent-to-sample ratio) was tested using 50 µL of each sample type: serum, plasma, NIST reference serum, and NIST reference plasma. **A)** Unconjugated bile acids, **B)** Glycine-conjugated bile acids, and **C)** Taurine-conjugated bile acids. This condition was evaluated without replication due to sample limitations.

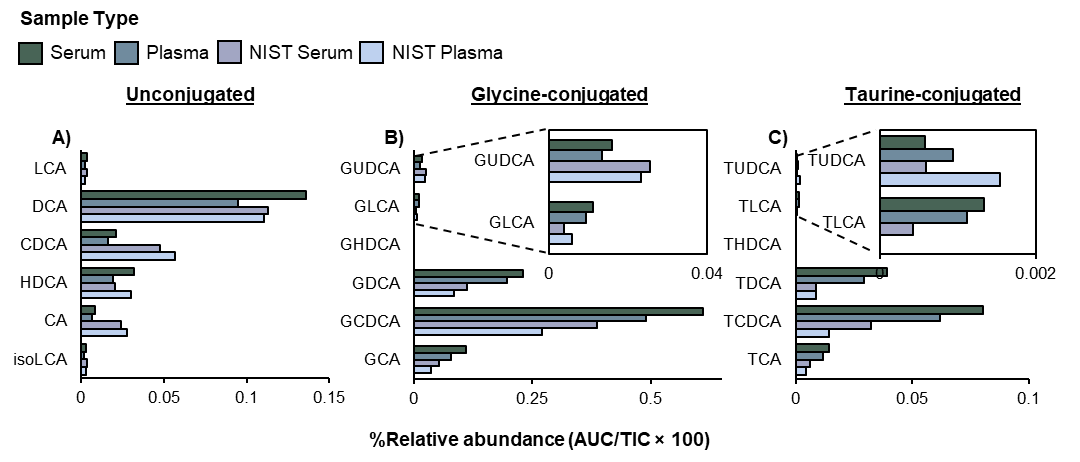

**Figure S5.** Representative collision cross section (CCS) distributions for bile acid features detected in positive-ion mode. CCS distributions are shown for common adducts, including **A)** [M+H]⁺, **B)** [M+Na]⁺, and **C)** [M+NH₄]⁺.

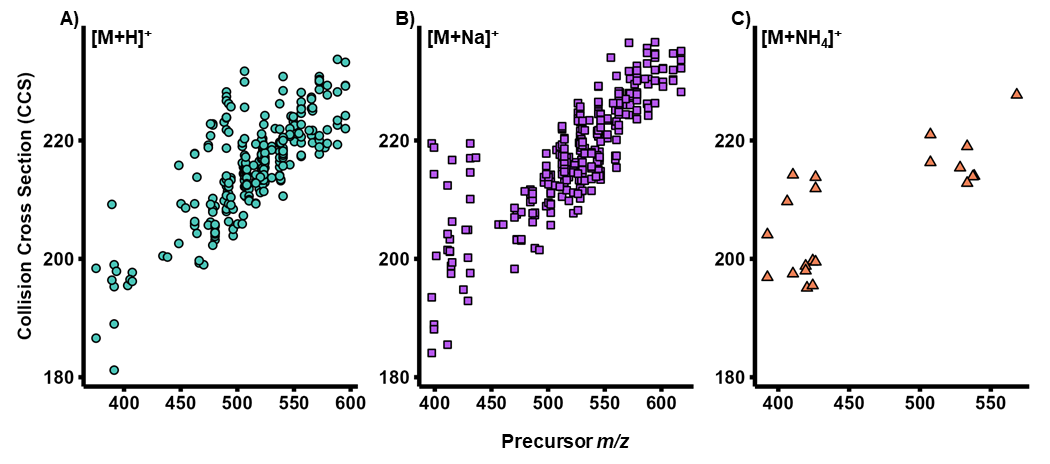
